# Fibroblast growth factor 21 regulates neuromuscular junction innervation through HDAC4 in denervation-induced skeletal muscle atrophy

**DOI:** 10.1101/2025.02.18.638653

**Authors:** Lirong Zheng, Takashi Sasaki, Liyang Ni, Yu Takahashi, Yoshio Yamauchi, Makoto Shimizu, Sato Ryuichiro

## Abstract

Skeletal muscles undergo atrophy in response to denervation and neuromuscular diseases. Understanding the mechanisms by which denervation drives muscle atrophy is crucial for developing therapies against neurogenic muscle atrophy. Here, we identify muscle-secreted fibroblast growth factor 21 (FGF21) as a key inducer of atrophy following muscle denervation. In denervated skeletal muscles, FGF21 is uniquely upregulated among the FGF family members and acts in an autocrine/paracrine manner to promote muscle atrophy. Silencing FGF21 in muscle prevents denervation-induced muscle wasting by preserving neuromuscular junction (NMJ) innervation. Conversely, forced expression of FGF21 in muscle reduces NMJ innervation, leading to muscle atrophy. Mechanistically, TGFB1 released by denervated fibro-adipogenic progenitors (FAPs) upregulates FGF21 through the JNK/c-Jun axis, which in turn reduces the cytoplasmic level of histone deacetylase 4 (HDAC4), culminating in muscle atrophy. HDAC4 knockdown abolishes the atrophy-resistant effects observed in FGF21-deficient denervated muscles, resulting in muscle atrophy. Our findings reveal a novel role and heretofore unrecognized mechanism of FGF21 in skeletal muscle atrophy, suggesting that inhibiting muscular FGF21 could be a promising strategy for mitigating skeletal muscle atrophy.

## Introduction

Skeletal muscle is a dynamic tissue essential for a wide variety of physiological processes. The maintenance of skeletal muscle mass relies on a delicate balance between protein synthesis and degradation (1). The innervation of skeletal muscle fibers by motor neurons is critical for preserving muscle size, structure, and function. Neuromuscular diseases, such as amyotrophic lateral sclerosis (ALS), are characterized by progressive motor neuron degeneration and significant loss of nerve supply to muscle fibers, leading to debilitating loss of muscle mass and function (neurogenic atrophy) and, in severe cases, death. Currently, there are no effective treatments for skeletal muscle loss in neurogenic disorders. Understanding the molecular mechanisms that regulate muscle mass is a prerequisite for developing novel therapeutics to combat muscle-wasting disorders.

Skeletal muscle-secreted cytokines, or myokines, play a pivotal role in enabling muscles to adapt to and influence systemic changes in response to external stimuli (2, 3). The fibroblast growth factor (FGF) family consists of 22 members, comprising hormone-like FGFs (FGF19, FGF21, and FGF23) and canonical FGFs. Unlike canonical FGFs, FGF21 lacks a heparin-binding domain, allowing it to circulate in the bloodstream as an endocrine factor (4–6). FGF21 is a myokine (7) that can also be secreted by other organs, such as the liver (8, 9), heart (10), and adipose tissues (11, 12). Research on FGF21 has predominantly focused on its beneficial effects on metabolic abnormalities, including obesity, type 2 diabetes, and non-alcoholic steatohepatitis (NASH), where it reduces fat mass and improves hyperglycemia, insulin resistance, and dyslipidemia (13–15). In skeletal muscle, FGF21 is secreted in response to the cellular stress pathways activated by exercise (16, 17), starvation (18), endoplasmic reticulum (ER) stress (19), autophagy impairment (20), mitochondrial dysfunction (20, 21), and aging (22, 23). While some studies report that FGF21 negatively impacts skeletal muscle mass and function (18, 24), its role in neurogenic skeletal muscle atrophy and the underlying mechanisms have not been investigated.

In this study, we demonstrate that skeletal muscles elevate FGF21 expression in response to denervation, promoting muscle atrophy in a non-endocrine manner. FGF21 acts on HDAC4 by reducing its cytoplasmic accumulation, leading to neuromuscular junction (NMJ) denervation. Conversely, the ablation of FGF21 enhances the cytoplasmic localization of HDAC4, enhancing NMJ reinnervation, thereby restoring muscle homeostasis. Moreover, the expression and function of FGF21 are regulated by TGFB1/JNK/c-Jun signaling derived from fibro-adipogenic progenitors (FAPs). Our findings reveal a novel role of FGF21 in neurogenic muscle atrophy by modulating NMJ innervation, positioning FGF21 as a potential therapeutic target for neuromuscular disorders.

## Results

### Fgf21 is upregulated in neurogenic atrophic skeletal muscles and is associated with muscle weakness

The degeneration of motor neurons occurs in various atrophic conditions, including aging (25), mitochondrial dysfunction (26), muscular dystrophy (27), neurogenic disorders (28), and myopathies (29). To identify potential key regulators associated with neurogenic muscle atrophy, we re-analyzed several public datasets (GSE18119, GSE48574, GSE52766, GSE87108, GSE49826) encompassing above mentioned atrophic models. Fgf21 expression was consistently elevated across all conditions (**Fig. 1A**) and significantly upregulated in our in-house models of atrophy induced by starvation, denervation, muscular dystrophy, and aging (**Fig. 1B**). After transecting sciatic nerve (**Fig. 1C**), severe muscle atrophy was induced as shown by decreased grip strength of mice, muscle size and weight, smaller myofiber size, and increased expression of pro-atrophy ubiquitin E3 ligases Fbxo32/Atrogin1 and Trim63/MuRF1 (**Fig. S1A-I**). Among FGF family members, only Fgf13 and Fgf21 were upregulated in response to denervation in the tibialis anterior (TA), with Fgf21 showing the most pronounced increase (**Fig. 1D**). Immunoblot and ELISA analyses confirmed elevated protein levels of FGF21 in denervated skeletal muscles (**Fig. 1E-G**). FGF21 signaling requires β-Klotho (Klb) and FGF receptors, and increased mRNA expressions of Klb, Fgfr1c, and Fgfr4 were observed in denervated muscles (**Fig. S1J**). Despite FGF21’s primary endocrine role, its plasma level remained unchanged following denervation (**Fig. 1H**). To exclude other FGF21-producing organs’ contributions, we assessed Fgf21 mRNA in the liver, epididymal white adipose tissue (eWAT), brain, and hindlimb muscles post-denervation and found that Fgf21 was exclusively upregulated in the denervated muscles (**Fig. S1K**). This muscle-specific increase was detectable from day 3 post-denervation and persisted until day 32 in the TA muscle (**Fig. S1L**). The Fgf21 response was corroborated in the denervated TA of rats from a public dataset (GSE201025, **Fig. S1M**). Analysis of the correlation between Fgf21 mRNA level, atrogene expression, muscle weight, size, and grip strength of mice revealed that Fgf21 expression was significantly induced by denervation and positively correlated with the expression of atrogenes upregulated by denervation (**Fig. 1I**). Conversely, these pro-atrophic factors showed a negative correlation with healthy muscle phenotype and function (**Fig. 1I**). Collectively, these findings suggest that endogenous Fgf21 expression may contribute to the progression of neurogenic muscle atrophy.

**Figure 1.**
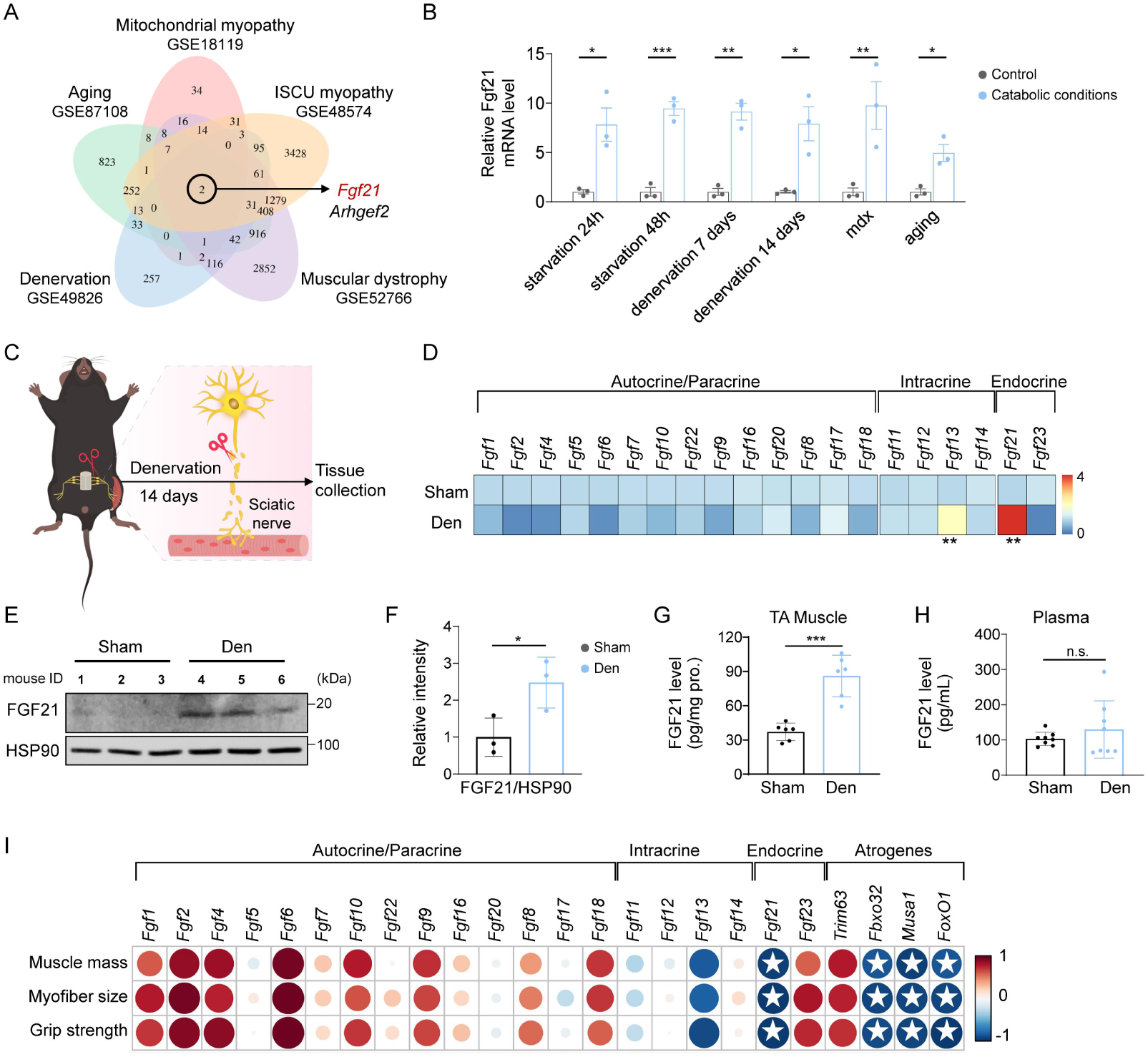
Fgf21 is upregulated in neurogenic atrophic skeletal muscles and is associated with muscle weakness. (A) Venn diagram of overlapping upregulated genes in five muscle atrophic conditions. (B) Fgf21 mRNA levels in the tibialis anterior (TA) muscle under starvation, denervation, dystrophy (mdx), and aging (n=3). (C) Schematic of denervation surgery (tissue harvested at 2 weeks). (D) Heatmap of mRNA levels of FGF members in denervated TA muscle (n=5). (E-F) Western blot and quantification of FGF21 protein in TA muscle (n=3). HSP90 was used as a loading control. (G) ELISA analysis of FGF21 protein levels in TA muscle (n = 6). (H) ELISA analysis of FGF21 protein levels in plasma (n = 8). (I) Correlation analysis between skeletal muscle phenotypes and mRNA levels of FGF family members and atrogenes (n=5).

### Fgf21 deficiency protects skeletal muscles from denervation-induced atrophy

To investigate the role of FGF21 in muscle atrophy, we used both loss-and gain-of-function approaches. Wild-type (WT) and Fgf21 knockout (Fgf21KO) mice underwent surgical denervation (**Fig. 2A, S2A**). As expected, both basal and denervation-induced Fgf21 expression was abolished in Fgf21KO mice (**Fig. S2B**). Grip strength normalized by body weight showed no significant difference between WT and Fgf21KO mice before denervation (**Fig. 2B**). Following denervation, both genotypes exhibited a decline in grip strength; however, Fgf21KO mice demonstrated a partial recovery compared to WT (**Fig. 2B**). Denervation reduced the muscle-to-body weight ratio in both genotypes, with no significant differences in normalized muscle weights between WT and Fgf21KO mice (**Fig. 2C**). The mean cross-sectional area (CSA) of sham-treated TA muscle was smaller in Fgf21KO mice compared to WT. However, post-denervation, Fgf21KO mice had a larger mean CSA and a greater proportion of large-sized fiber distribution than WT (**Fig. 2D, E; S2C**). We evaluated protein degradation to understand the preservation of muscle size and strength in Fgf21KO mice and found that protein degradation markers (Atrogin1, MuRF1, LC3II) were upregulated by denervation but suppressed in Fgf21KO mice (**Fig. 2F, G**).

**Figure 2.**
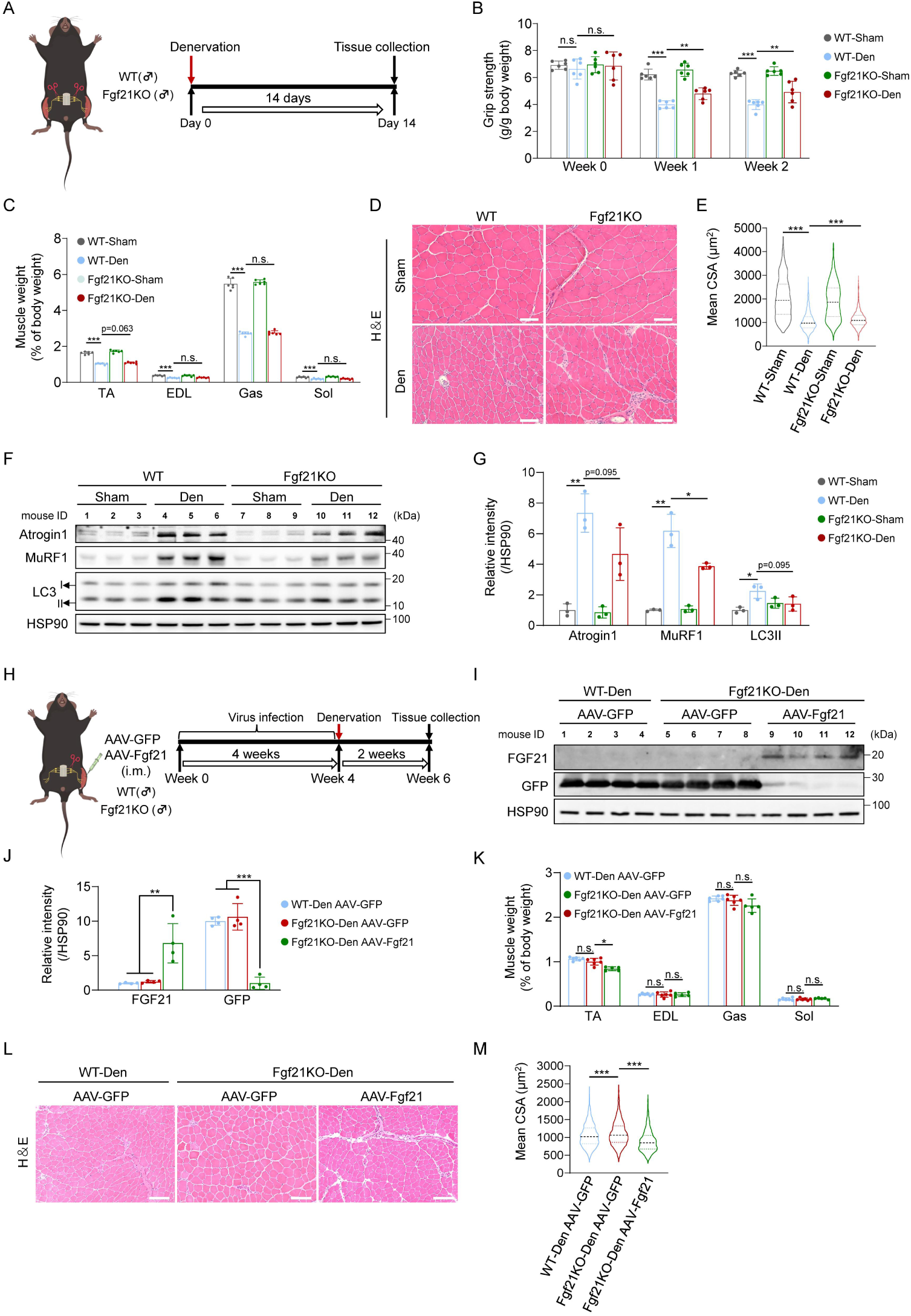
Fgf21 deficiency protects skeletal muscles from denervation-induced atrophy. (A) Experimental timeline of 2-week denervation in WT and Fgf21KO mice. (B) Four-limb grip strength (g/g body weight) of mice (n = 6). (C) The ratio of limb muscle weight to body weight (n = 6). (D-E) Left panels: Representative cross-sectional images of TA muscle. Scale bar, 50 μm. Right panels: Quantification of mean cross-sectional area (CSA) (n = 4). (F-G) Western blot and quantification of protein degradation markers in TA muscle (n=3). HSP90 was used as a loading control. (H) Rescue experiment design: TA muscle of WT/Fgf21KO mice injected with GFP-or Fgf21-expressing virus prior to 2-week denervation. (I-J) Western blot and quantification of FGF21 protein levels in TA muscle (n=4). HSP90 was used as a loading control. (K) The ratio of limb muscle weight to body weight post-rescue (n=5-6). (L-M) Left panels: Representative cross-sectional images of TA muscle. Scale bar, 50 μm. Right panels: Quantification of mean CSA (n = 6).

Given Fgf21 deficiency’s protective effect against denervation-induced atrophy, we examined the impact of Fgf21 overexpression in TA muscle. AAV-mediated Fgf21 overexpression in TA muscle followed by denervation significantly elevated circulating FGF21 level, irrespective of denervation status (**Fig. S3A-E**). Fgf21 overexpression reduced grip strength and relative TA muscle weight post-denervation (**Fig. S3F, G**). Fgf21 overexpression alone decreased myofiber size, with a smaller CSA distribution, and exacerbated denervation-induced reductions in myofiber size (**Fig. S3H-J**).

To confirm the intramuscular role of Fgf21, rescue experiments were conducted by AAV-mediated Fgf21 overexpression in Fgf21-null TA muscles, followed by denervation (**Fig. 2H-J**). Plasma FGF21 was undetectable in Fgf21KO mice, but Fgf21 overexpression significantly increased FGF21 plasma levels (**Fig. S3K**). TA muscle weight was similar between GFP-transduced WT and Fgf21KO mice; however, Fgf21 overexpression in Fgf21-ablated TA muscles reduced TA muscle weight (**Fig. 2K**). The mean CSA was larger in Fgf21KO denervated mice compared to WT, but Fgf21 overexpression in Fgf21KO denervated mice nullified this effect, reducing myofiber size and shifting the distribution towards smaller sizes (**Fig. 2L, M; S3L**). These data confirm that Fgf21 induces skeletal muscle atrophy, and its ablation protects muscle against denervation-induced atrophy.

### Fgf21 deficiency improves NMJ innervation upon denervation

To understand how Fgf21 deficiency resists denervation-induced muscle atrophy, we performed RNA-Seq on TA muscle from WT and Fgf21KO mice at two weeks post-denervation. Heatmap showed altered gene expression patterns due to denervation and/or Fgf21KO (**Fig. S4A**). A Venn diagram displayed common genes among different comparisons (**Fig. S4B**). Differentially expressed gene (DEG) analysis (adjusted p-value < 0.05, absolute fold change > 1.5) revealed numerous upregulated and downregulated genes (**Fig. S4C**). KEGG pathway enrichment analysis indicated that denervation-and Fgf21-dependent gene changes were associated with Axon guidance, Neurotrophin signaling, Cytokine-cytokine receptor interaction, and Calcium signaling pathways (**Fig. S4D-G**). Particularly, Axon guidance is crucial for skeletal muscle function, highlighting the neuromuscular junction (NMJ) as a critical structure.

NMJ damage is common in muscle atrophy models, and we assessed the NMJ integrity in denervated and non-denervated TA muscles from WT and Fgf21KO mice. WT mice exhibited reduced NMJ innervation post-denervation, whereas Fgf21KO mice showed partial restoration (**Fig. 3A, B**). NMJs remained innervated in sham-operated muscles of both genotypes (**Fig. 3A, B**). Denervation reduced NMJ structural proteins NF-M and SV2 in both genotypes compared to sham controls, but Fgf21KO mice showed partial restoration (**Fig. 3C, D**). No significant differences in NMJ protein levels were observed between sham-operated WT and Fgf21KO mice (**Fig. 3C, D**). In vitro, administration of recombinant FGF21 protein in muscle cells significantly reduced the expression of NMJ-associated genes (AChRs, Lrp4, Musk, App, and Dok7), and post-synaptic AChR intensity along myotube surface (**Fig. 3E-G**).

**Figure 3.**
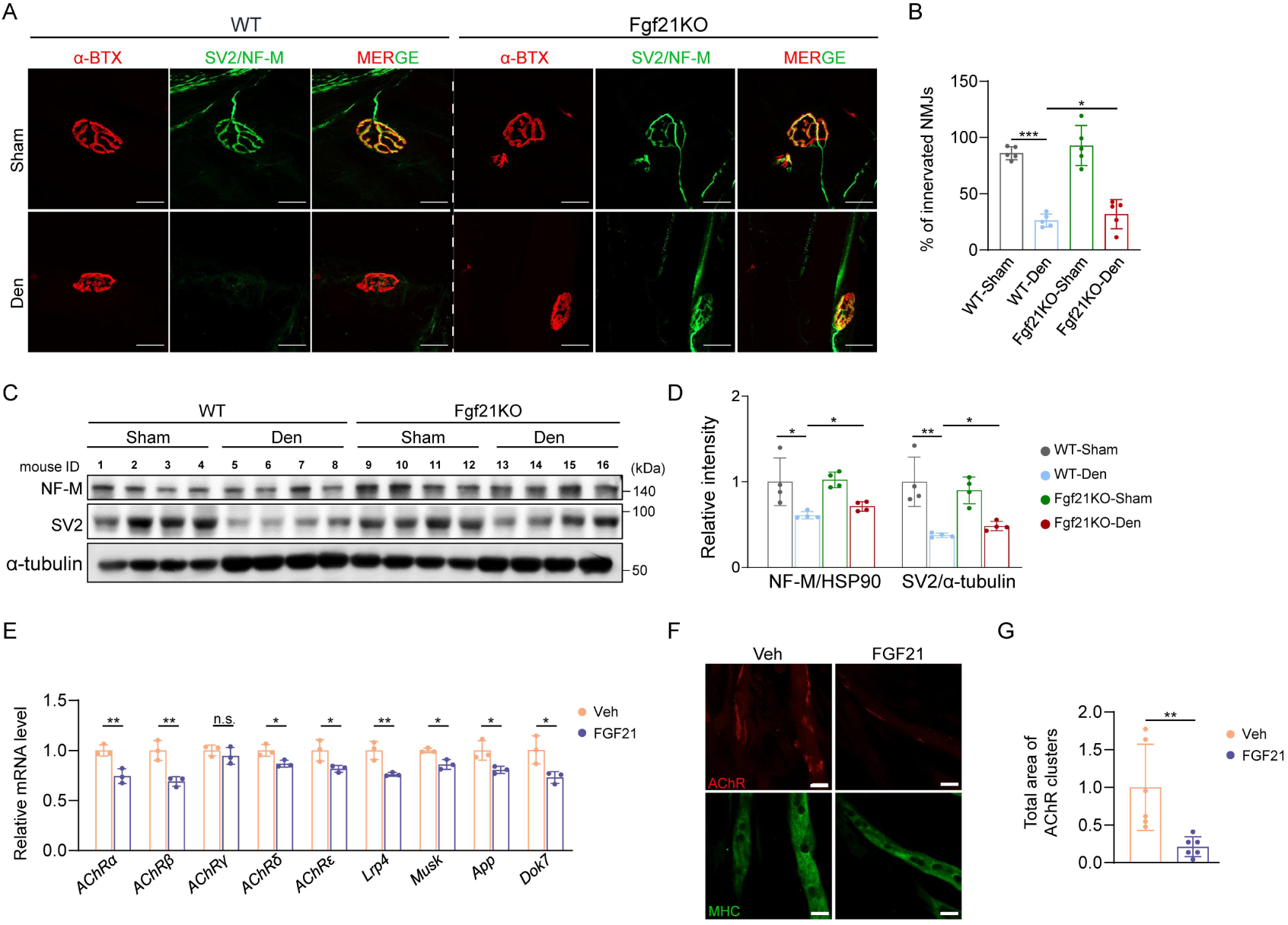
Fgf21 deficiency improves NMJ innervation. (A-D) WT and Fgf21KO mice were subjected to denervation for 2 weeks. (A, B) Representative images and quantification of neuromuscular junction (NMJ) innervation in the extensor digitorum longus (EDL) muscle (n = 5). (C-D) Western blot and quantification of NF-M and SV2 proteins levels in EDL muscle (n=4). HSP90 and α-tubulin were used as loading controls. (E-F) C2C12 myotubes were treated with 1 μg/mL recombinant FGF21 protein for 24 hours. (E) mRNA levels of NMJ-related genes (n=3). (F) Representative images and quantification of AChR intensity in C2C12 myotubes. Scale bar, 20 μm.

These findings indicate that the muscle atrophy-resisting effect of Fgf21 deficiency is associated with improved NMJ innervation.

### Fgf21 deficiency resists muscle atrophy via HDAC4

Class IIa histone deacetylases (HDACs), particularly HDAC4, play a crucial role in neural activity and muscle responses to denervation. To explore the mechanisms underlying improved NMJ reinnervation, we investigated the relationship between FGF21 and HDAC4. The mRNA level of HDAC4 was increased following denervation and was unaffected by Fgf21 deficiency (**Fig. 4A**). However, HDAC4 protein level was further elevated in Fgf21KO mice post-denervation, suggesting a post-transcriptional regulation of HDAC4 (**Fig. 4B, C**). Other Class IIa HDACs, HDAC5, and HDAC7 were upregulated by denervation but unaffected by Fgf21 deficiency (**Fig. S5A, B**).

**Figure 4.**
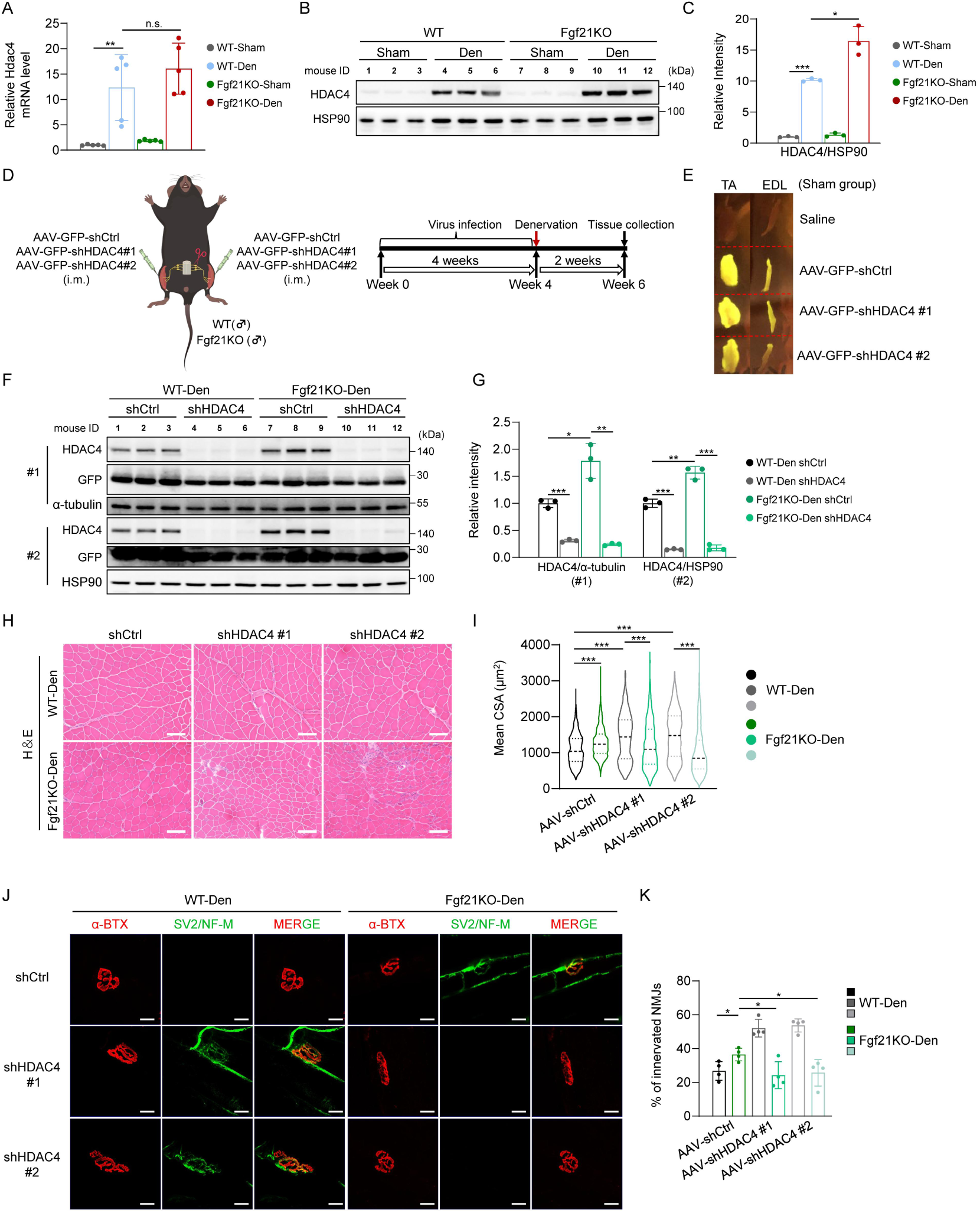
Fgf21 deficiency resists muscle atrophy via HDAC4. (A) Fgf21 mRNA levels in denervated TA muscle of WT and Fgf21KO mice (n=5). (B-C) Western blot and quantification of HDAC4 protein levels in denervated TA muscle (n=3). HSP90 was used as loading controls. (D) Experimental design of HDAC4 knockdown via AAV-shRNA. (E) Representative images of AAV-transduced GFP expression in TA and EDL muscles. (F-G) Western blot and quantification of HDAC4 knockdown efficiency in denervated TA muscle (n=3). α-tubulin/HSP90 loading controls were used as loading controls. (H, I) Left panels: Representative cross-sectional images of the denervated TA muscle. Scale bar, 50 μm. Right panels: Quantification of mean CSA (n = 5). (J, K) Representative images and quantification of NMJ innervation in the denervated EDL muscle of mice (n = 4).

We then took advantage of AAV-mediated delivery of shRNA targeting mouse Hdac4 in TA muscle post-denervation to examine the functional implications of HDAC4 (**Fig. 4D**). Successful delivery of the virus was confirmed by GFP expression (**Fig. 4E**), and HDAC4 knockdown efficiency was validated at protein level (**Fig. 4F, G; S5C, D**). In non-denervated TA muscles, both Fgf21 deficiency and Hdac4 knockdown decreased mean CSA compared to control shRNA, highlighting their roles under physiological conditions (**Fig. S5E-G**). Two weeks post-denervation, WT muscles injected with control shRNA showed a 50% CSA reduction compared to sham-operated muscles (**Fig. 4H, I; S9H**), while Hdac4 knockdown in WT denervated muscles increased CSA compared to controls (**Fig. 4H, I; S5H**). In contrast, Hdac4 knockdown in Fgf21KO denervated muscles further reduced CSA compared to control shRNA, indicating a context-dependent role of HDAC4 in muscle atrophy (**Fig. 4H, I; S5H**). NMJ innervation was restored by either Fgf21 deficiency or Hdac4 knockdown in WT-denervated muscles but was lost when Hdac4 was knocked down in Fgf21-null muscles (**Fig. 4J, K**). The NMJ innervation was not affected by either Fgf21 deficiency or Hdac4 knockdown in sham condition (**Fig. S5I, J**).

To understand the role of HDAC4 in neurogenic muscle atrophy in the context of Fgf21 deficiency, we examined its phosphorylation-regulated subcellular localization. Denervation upregulated cytoplasmic and nuclear HDAC4 levels in WT muscles (**Fig. S6A, B**), while Fgf21 deficiency further increased cytoplasmic HDAC4 level, with no effect on nuclear HDAC4 level (**Fig. S6A, B**). Corresponding increases in HDAC4 phosphorylation at two serine sites were observed (**Fig. S6C, D**). AMPK, a kinase that regulates muscle atrophy, showed elevated phosphorylation levels in WT-denervated muscles and further increases in Fgf21KO muscles, indicating the involvement of AMPK regulation of HDAC4 phosphorylation (**Fig. S6E-G**). To test the hypothesis, we treated C2C12 cells with AICAR (an AMPK activator) and/or recombinant FGF21 protein. AICAR increased the cytoplasmic localization of HDAC4, an effect that was reversed by FGF21 addition, leading to increased nuclear HDAC4 localization (**Fig. S6H-K**). In addition, AICAR-induced phosphorylation of AMPK and HDAC4 was suppressed by FGF21, indicating that FGF21 inhibits the AMPK-mediated cytoplasmic location of HDAC4 (**Fig. S6L, M**). In vivo, AAV-mediated FGF21 overexpression reduced cytoplasmic HDAC4 accumulation without affecting its nuclear level (**Fig. S6N-Q**), confirming the regulation of FGF21 on AMPK-mediated HDAC4 localization. These data demonstrate that AMPK-mediated cytoplasmic retention of HDAC4 is crucial for the muscle atrophy-resisting effects observed in Fgf21 deficiency.

Interestingly, as Fgf21 is a key downstream factor of ER stress, we also observed altered calcium signaling in denervated Fgf21KO compared to WT muscles (**Fig. S4E**), providing a hint of Fgf21’s involvement in denervation-induced ER stress. Recombinant FGF21 protein treatment caused elevated ER stress levels in C2C12 cells (**Fig. S7A, B**). When C2C12 myotubes were treated with chemical ER stress inducers Tunicamycin (Tm) or Thapsigargin (TG), elevated ER stress levels were accompanied by decreased AChR intensity along the myotube surface (**Fig. S7C-E**), indicating a destructive effect of ER stress on NMJ integrity. To consolidate the idea, we silenced Fgf21 before Tm or Tg treatment in C2C12 cells. Upregulated ER stress levels by Tm or Tg were significantly downregulated after Fgf21 silencing (except for Perk), along with recovered AChR intensity (**Fig. S7F-H**), suggesting the role of Fgf21 in mediating ER stress-caused NMJ destruction. Moreover, Fgf21 deficiency alleviated denervation-induced ER stress in TA muscles, whereas Hdac4 knockdown in Fgf21-null muscles negated Fgf21 deficiency’s protective effects, exacerbating ER stress (**Fig. S7I**), stressing the potential contribution of FGF21-HDAC4 axis-mediated ER stress in neurogenic muscle atrophy.

### TGFB1 promotes myotube atrophy via FGF21-mediated NMJ damage

We then investigated how Fgf21 is regulated during denervation-induced muscle atrophy. RNA-seq data from denervated TA muscles of WT mice showed enriched TGFB and MAPK signaling pathways, suggesting a link between TGFB signaling and FGF21 expression (**Fig. S4D**). Denervation significantly increased TGFB1 protein level in TA muscle (**Fig. 5A, B; S8A**) and upregulated Tgfb1 mRNA specifically in skeletal muscles (**Fig. 5C**), while plasma TGF-β1 level remained unchanged (**Fig. S8B**). In C2C12 myotubes, TGFB1 treatment elevated FGF21 expression at both mRNA and protein levels, as well as its secretion into the culture medium (**Fig. 5D-F; S8C**). TGFB1 also induced muscle atrophy, evidenced by decreased myotube diameter, reduced MHC protein level, and increased expression of pro-atrophy markers Atrogin1 and MuRF1 (**Fig. S8D-G**). Similarly, FGF21 treatment promoted muscle atrophy in C2C12 myotubes (**Fig. S8H-K**).

**Figure 5.**
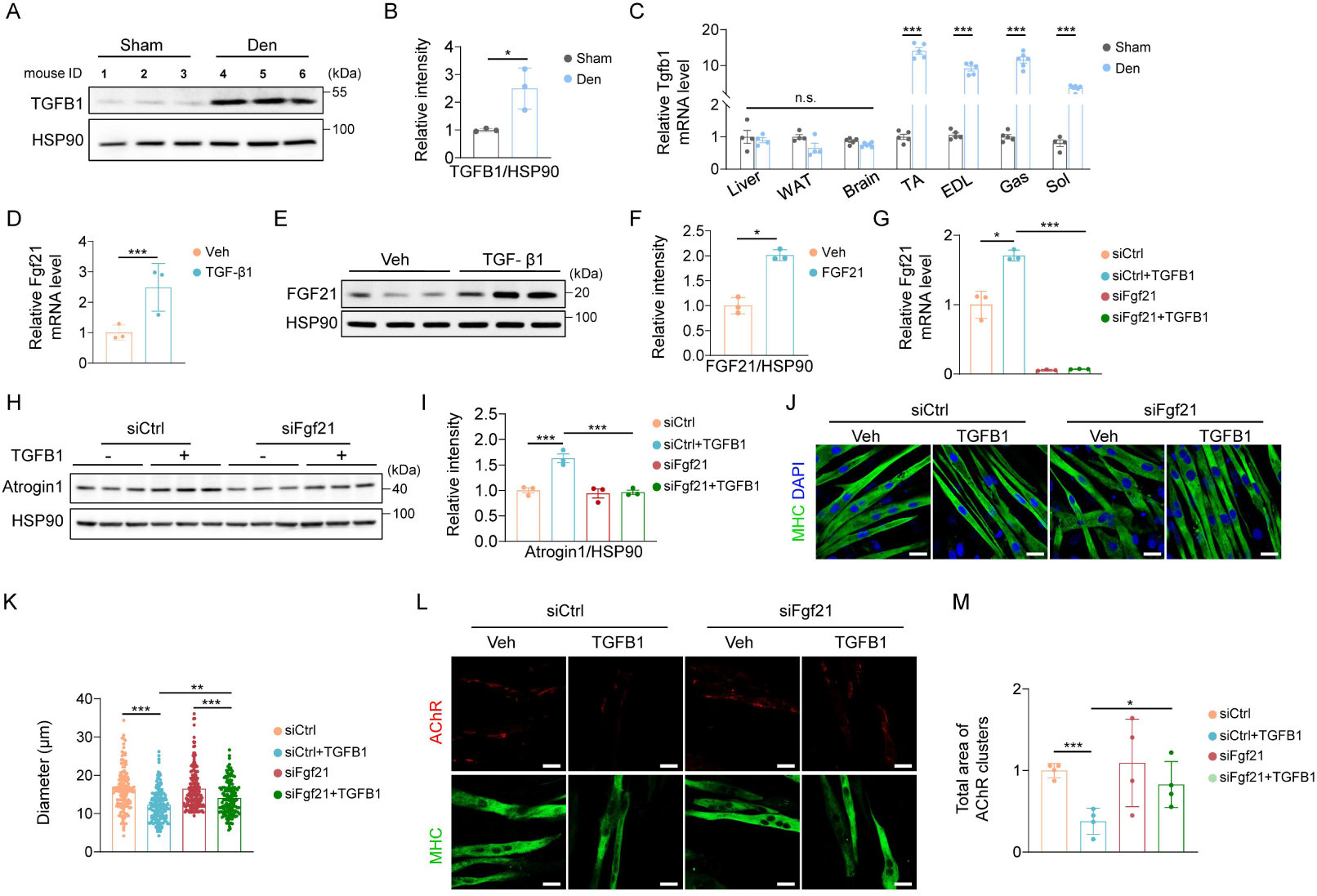
TGFB1 promotes myotube atrophy via FGF21-mediated NMJ damage. (A-B) Western blot and quantification of TGFB1 protein levels in 2-week denervated TA muscle (n=3). HSP90 loading control was used as a loading control. (C) Tgfb1 mRNA levels across tissues post-denervation (n=4-6). (D-F) TGFB1 (5 ng/mL) effects on FGF21 in C2C12 myotubes (n=3): (D) Fgf21 mRNA levels (4 h). (E-F) Western blot and quantification of TGFB1 protein levels (24 h). HSP90 loading control was used as a loading control. (G-P) Fgf21 siRNA knockdown in TGFB1-treated myotubes (n=3): (G) Fgf21 mRNA levels (4 h). (H-I) Western blot and quantification of Atrogin1 protein levels (24 h). HSP90 was used as a loading control. (J-K) Representative images and quantification of myotube diameter (24 h). (L-M) Representative images and quantification of AChR intensity (24 h). Scale bar, 20 μm.

To determine if elevated FGF21 contributes to TGFB1-induced atrophy, Fgf21 was silenced in C2C12 cells before TGF-β1 treatment. Silencing Fgf21 blocked both basal and TGFB1-induced upregulation of Fgf21 (**Fig. 5G**) and suppressed TGFB1-induced Atrogin1 expression, restoring myotube diameter (**Fig. 5H-K**). Under physiological conditions, Fgf21 silencing had no effect in the absence of TGFB1 treatment (**Fig. 5H-K**). Interestingly, Fgf21 mRNA was efficiently induced only in mature myotubes, not in undifferentiated myoblasts (**Fig. S8L**), indicating that TGFB1-induced muscle atrophy via FGF21 occurs specifically in muscle fibers.

We also investigated if TGFB1 induces NMJ damage through Fgf21. TGFB1 treatment reduced both NMJ-related gene expression and AChR intensity (**Fig. S8D-F**). Silencing Fgf21 before TGFB1 treatment partially restored AChR intensity (**Fig. 5L, M**). These findings suggest that TGFB1 increases Fgf21 to cause NMJ damage, contributing to muscle atrophy.

### TGFB1 regulates FGF21 expression through non-canonical JNK/c-Jun axis

We next investigated how TGFB1 regulates Fgf21 expression during muscle atrophy. TGFB signaling involves the canonical Smad pathway and non-canonical kinase pathways (**Fig. S9A**). To explore the mechanism of TGFB1-induced Fgf21 upregulation, we first examined the Smad pathway. C2C12 myotubes were pre-treated with SB-525334 (a TGFB type I receptor inhibitor) or SIS3 (a Smad3 phosphorylation inhibitor) before TGFB1 exposure. TGFB1 increased the expression of Smad target genes Pai1 and Ctgf, the effect that was abolished by SB-525334 and partially attenuated by SIS3 (**Fig. S9B**). Although TGFB1 also upregulated Fgf21, only SB-525334 partially mitigated the increase, while SIS3 had no effect (**Fig. S9B**). Additionally, Smad3 silencing by siRNA reduced the TGFB1-induced upregulation of Pai1 and Ctgf but did not affect Fgf21 level, indicating that Fgf21 is not regulated by canonical Smad signaling (**Fig. S9C-E**).

Next, we explored non-canonical pathways. C2C12 myotubes were pre-treated with various kinase inhibitors before TGFB1 treatment (**Fig. S9F-M**). We found that TGFB1 elevated JNK-targeted c-Jun phosphorylation, which was downregulated by both SB-525334 and the JNK inhibitor SP600125 (**Fig. 6A, B**). Co-treatment with TGFB1 and either SB-525334 or SP600125 reduced TGFB1-induced c-Jun, c-Fos, and Fgf21 expression (**Fig. 6C**). In addition, pre-treatment with SP600125 followed by Anisomycin (a JNK activator) enhanced JNK and c-Jun phosphorylation, which was suppressed by SP600125 (**Fig. 6D, E**). Anisomycin also upregulated JNK target genes and Fgf21 expression, effects that were reduced by SP600125 (**Fig. 6F**). JNK and c-Jun protein levels were elevated in WT denervated TA muscle compared to non-denervated muscle (**Fig. S9N, O**). Moreover, pre-treatment with SP600125 followed by TGFB1 in C2C12 myotubes partially restored myotube diameter and reduced Atrogin1 expression (**Fig. 6G-J**). These findings demonstrate that TGFB1 regulates Fgf21 expression through the non-canonical JNK/c-Jun axis, rather than the canonical Smad pathway, to promote muscle atrophy.

**Figure 6.**
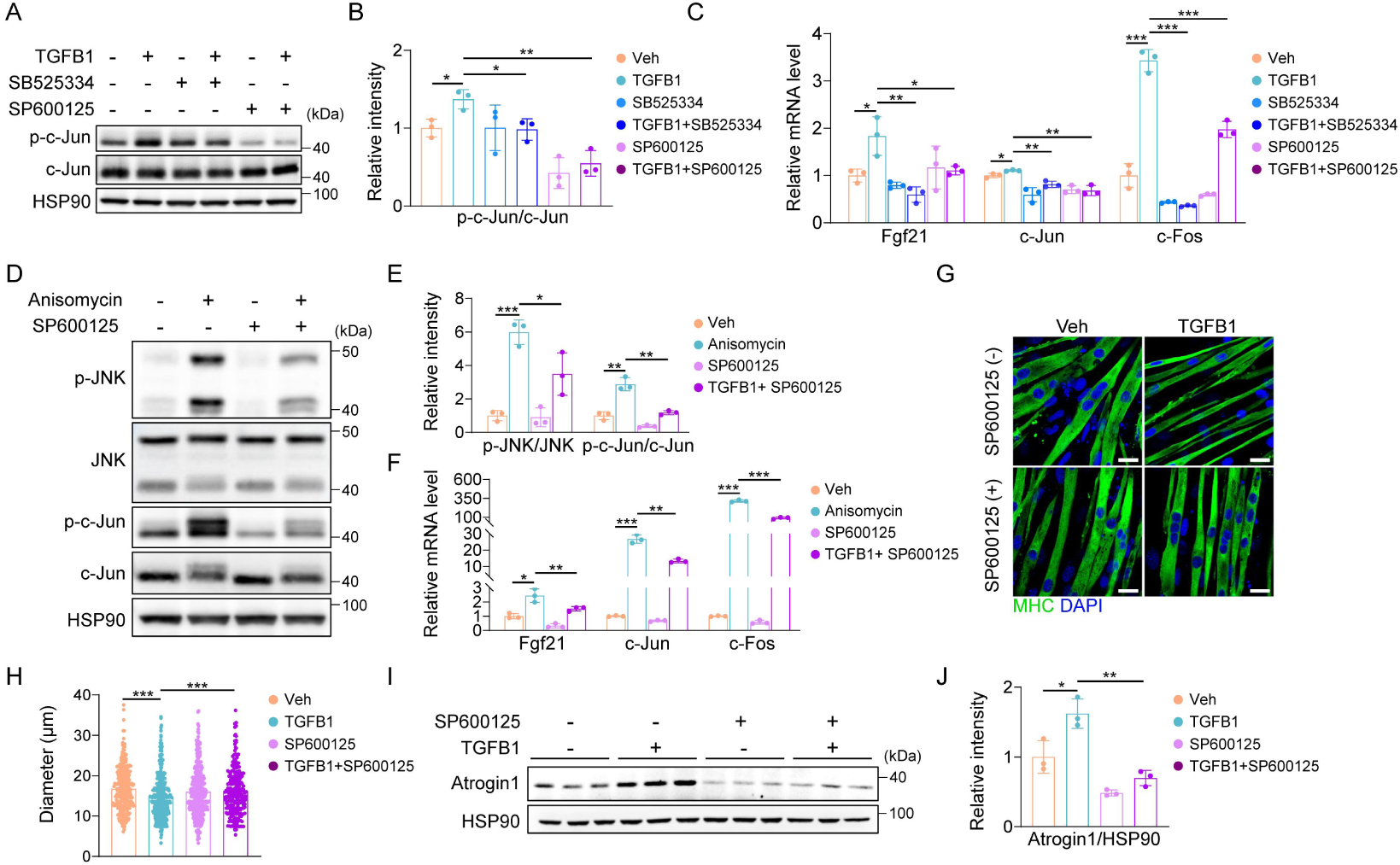
TGFB1 regulates FGF21 expression through non-canonical JNK/c-Jun axis. (A-C) In C2C12 myotubes treated with TGFB1 ± inhibitors (n = 3): (A, B) Western blot and quantification of phospho-and total c-Jun protein levels (1 h). HSP90 was used as a loading control. (C) mRNA levels of Fgf21, c-Jun, and c-Fos (4 h). (D-F) In C2C12 myotubes treated with Anisomycin ± inhibitors (n = 3): (D, E) Western blot and quantification of phospho-and total JNK and c-Jun protein levels (30 min). HSP90 was used as a loading control. (F) mRNA levels of Fgf21, c-Jun, and c-Fos (1 h). (G-J) In C2C12 myotubes treated with TGFB1 ± SP600125 (JNK inhibitor) (n = 3): (G, H) Representative images and quantification of myotube diameter (48 h). Scale bar, 20 μm. (I, J) Western blot and quantification of Atrogin1 protein levels (48 h). HSP90 was used as a loading control.

### Denervation-activated FAPs secrete TGFB1 to promote muscle atrophy

To understand the source of denervation-upregulated TGFB1, we focus on fibro-adipogenic progenitors (FAPs), which are known to secrete Activin A (a member belonging to the TGFB superfamily) to promote muscle atrophy. FAPs from denervated muscles (Den-FAPs) were isolated using Magnetic-activated cell sorting (MACS) and compared with those from sham-operated muscles (Sham-FAPs) (**Fig. 7A; S10A**). The number of FAPs was increased in denervated muscles compared to sham muscles (**Fig. S10B**). TGFB1 expression was significantly higher in Den-FAPs at both mRNA and protein levels (**Fig. 7B, C; S10C**), and Den-FAPs-derived conditioned medium (Den-CM) showed higher TGFB1 level than Sham-CM (**Fig. 7D**). Treating C2C12 myotubes with Den-CM reduced myotube diameter and increased Atrogin1 protein level (**Fig. S10D-G**), effects that were reversed by TGFB inhibitor SB525334 (**Fig. 7E-H**). These data indicate that TGFB1 released from FAPs contributes to muscle atrophy.

**Figure 7.**
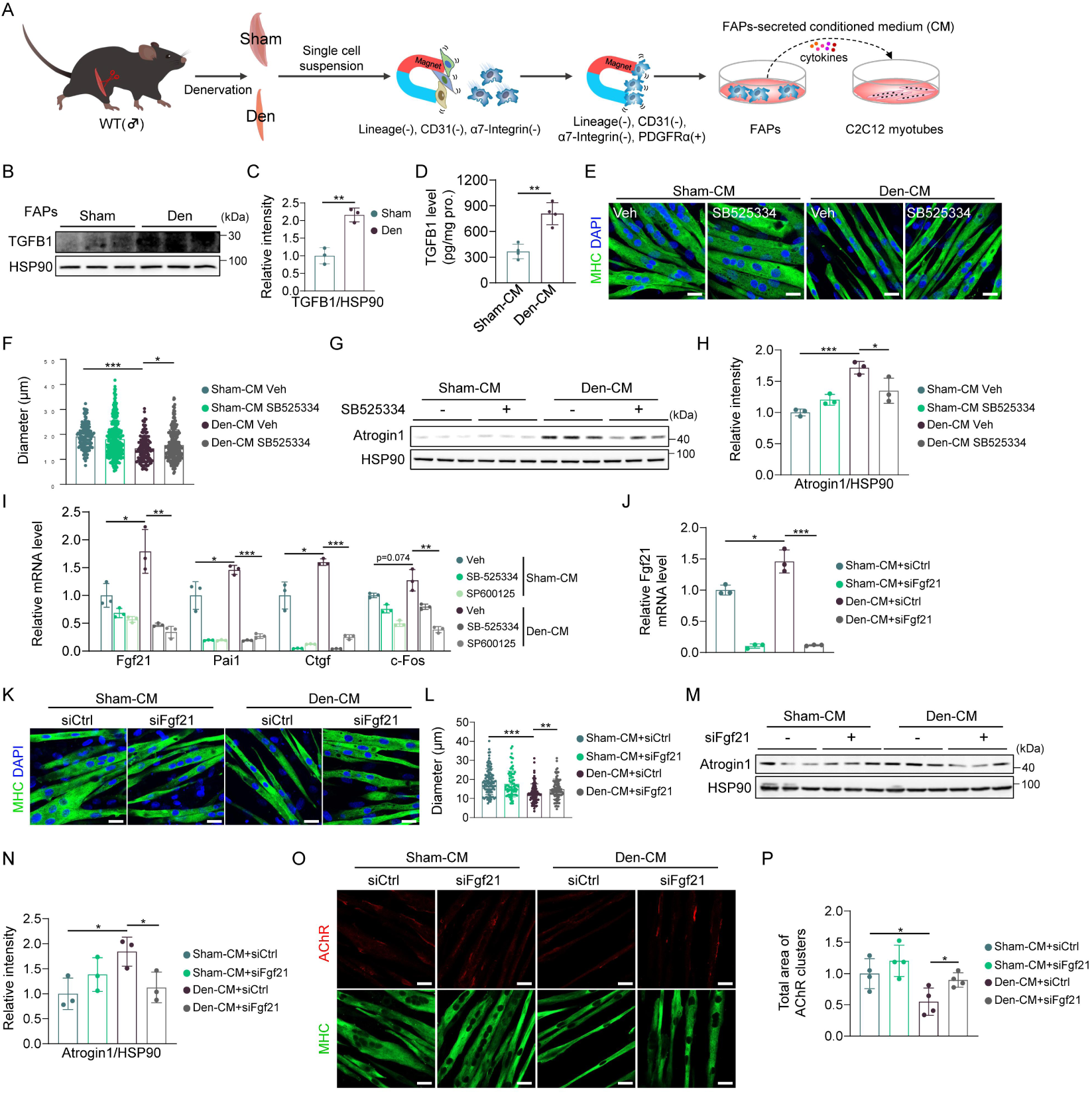
Denervation-activated FAPs secrete TGFB1 to promote muscle atrophy. (A) Schematic diagram of FAPs isolation and conditioned medium (CM) collection. (B, C) Western blot and quantification of TGFB1 protein levels in FAPs (n = 3). HSP90 was used as a loading control. (D) ELISA analysis of TGFB1 protein levels in CM (n = 4). (E–I) C2C12 myotubes were pre-treated with 10 µM SB525334 and/or 10 µM SP600125 for 30 minutes, followed by treatment with CM for the indicated times (n = 3). (E, F) Representative images and quantification of myotube diameter (48 h). Scale bar, 20 μm. (G, H) Western blot and quantification of Atrogin1 protein levels (48 h). HSP90 was used as a loading control. (I) mRNA levels of Fgf21, Pai1, c-Jun, and c-Fos (24 h). (J–P) C2C12 myotubes were transfected with siFgf21 and subsequently treated with CM for 48 hours (n = 3). (J) mRNA levels of Fgf21. (K, L) Representative images and quantification of myotube diameter. Scale bar, 20 μm. (M, N) Western blot and quantification of Atrogin1 protein levels. HSP90 was used as a loading control. (O, P) Representative images and quantification of AChR intensity. Scale bar, 20 μm.

Treating C2C12 myotubes with Den-CM increased the expressions of TGFB target genes, including Pai1, Ctgf, c-Fos (which tended to increase), and Fgf21, all of which were suppressed by SB-525334 and SP600125 (**Fig. 7I**). Silencing Fgf21 mitigated Den-CM-induced myotube atrophy and the increase in Atrogin1 protein level (**Fig. 7J-N**). Furthermore, to determine if TGFB1 released by FAPs could provoke NMJ damage, we treated C2C12 myotubes with Den-CM. Den-CM reduced AChR intensity compared to Sham-CM in C2C12 myotubes (**Fig. S10H, I**). To investigate if TGFB1 released by FAPs induces NMJ damage via Fgf21, Fgf21 was silenced in C2C12 cells before exposure to Den-CM. Fgf21 knockdown restored AChR intensity, which was downregulated by Den-CM alone (**Fig. 7O-P**), suggesting TGFB1 secreted by Den-FAPs promotes atrophy through Fgf21-mediated NMJ damage.

All these data combined to demonstrate that FAPs are activated upon denervation to release TGFB1, which induces the expression of Fgf21 in myofibers, leading to muscle atrophy.

## Discussion

Since its discovery in 2000 (30), FGF21 has been recognized as a stress-induced hormone synthesized by various metabolic organs and has emerged as a novel therapeutic target for treating obesity and related metabolic syndromes (5, 31, 32). A study of metabolic dysfunction-associated steatohepatitis (MASH) has reported that muscle-directed AAV1-FGF21 gene therapy reversed MASH progression (33). Despite its benefits against metabolic disorders, the role of FGF21 in skeletal muscle remains largely unexplored.

FGF21 is induced in myogenic cell lines during differentiation (34–36) and promotes myogenesis (36). It enhances glucose uptake in human primary myotubes (37), improves mitochondrial function (38), and protects against inflammation-mediated muscle atrophy in murine myotubes (39), implicating FGF21 as a positive regulator of skeletal muscle maturation and energy metabolism. In contrast, direct expression of FGF21 in skeletal muscle induces atrophy through mitophagy activation (18), and pharmacological administration of FGF21 leads to significant skeletal muscle loss via increased HPA axis activity (24), indicating FGF21 negatively affects skeletal muscle via different signaling pathways. Elevated circulating levels of FGF21 have been observed in patients with neuromuscular disorders (40), mitochondrial myopathy (41), and sarcopenia (42), suggesting that the pathophysiological role of FGF21 in skeletal muscle may be context-specific.

Our study reveals a critical role for FGF21 in regulating neurogenic muscle atrophy. We found that FGF21 expression is inversely correlated with muscle mass and function following denervation. FGF21 deficiency significantly alleviates muscle atrophy following denervation by promoting the cytoplasmic accumulation of HDAC4, which enhances NMJ reinnervation, thereby contributing to muscle homeostasis. The upregulation of FGF21 in response to denervation is driven by TGFB1-mediated JNK/c-Jun signaling originating from muscle-resident FAPs. The autocrine/paracrine action of FGF21 in muscle-specific contexts, as observed in our study, may be due to the consequences of sciatic nerve transection, which predominantly impacts nerve-controlled limb muscles. Additionally, skeletal muscles, which primarily consist of structural proteins and glycogen, align with FGF21’s role in promoting autophagic degradation, allowing atrophic muscles to meet their energy demands more efficiently without mobilizing resources from distant organs.

Neuromuscular junctions (NMJs), the synaptic interfaces between motor neuron branches and muscle cells, consist of three elements: the pre-synaptic motor nerve terminal, the intrasynaptic synaptic basal lamina, and the post-synaptic muscle fiber and muscle membrane (43). The NMJ is a specialized structure for the transmission of action potential from motor neurons to muscle fibers to initiate muscle contraction. FGF21 has been reported to function in the central nervous system to regulate memory (44), determine diet preference (45), and attenuate neurodegeneration (46). On the other hand, FGF21 also plays controversial roles in the peripheral nervous system. One group has reported that FGF21 facilitates peripheral nerve regeneration after injury (47), while another study reveals that FGF21 acts as a negative regulator for myelin development in Schwann cells (48). Here, we found that pharmacological administration of recombinant FGF21 protein suppresses AChR expression in muscle cells, leading to muscle atrophy. It seems that the function of FGF21 is related to the cell stage, with its roles likely being opposite in the proliferative state compared to the terminal differentiated state, which warrants further investigation. Of note, the protective effect of FGF21 against muscle atrophy is mediated by the crucial kinase AMPK, a well-established regulator of autophagic and catabolic processes (49). Our finding that AMPK activation contributes to NMJ improvement aligns with recent reports, which suggest that AMPK mediates the maintenance and plasticity of motoneurons and the NMJ, and genetic, pharmacological, or physiological activation of AMPK induces beneficial phenotypic remodeling in neuromuscular disorders (NMDs) (50).

FGF21/AMPK axis mediates its effects by stimulating the phosphorylation of HDAC4, a Class IIa histone deacetylase, resulting in its cytoplasmic retention. HDAC4 is known to regulate muscle development, differentiation, and fiber-specific transcriptional programs (51), but its role in neurogenic muscle atrophy remains controversial. Moresi et al. report that HDAC4 promotes muscle atrophy upon denervation by activating myogenin-mediated proteolysis (52). Williams et al. report that miR-206 slows ALS progression by inhibiting HDAC4, thereby promoting neuromuscular synapse regeneration (53). On the contrary, Pigna et al. show that HDAC4 preserves skeletal muscle structure following long-term denervation (54), and that deletion of HDAC4 induces earlier onset of ALS (55). However, to date, no study has specifically reported the role of HDAC4’s distinct subcellular localization in denervated skeletal muscles. Nuclear HDAC4 has long been recognized as a transcriptional repressor forming a complex with N-CoR/SMRT/HDAC3 to inhibit MEF2-dependent gene transcription (56, 57). Regarding cytoplasmic HDAC4, it’s reported to either prevent ER stress-induced apoptotic cell death (58) and mediate muscle repair in muscular dystrophy (59) or promote muscle atrophy via deacetylation of cytosolic substrates (60). Our study shows that knocking down cytoplasmic HDAC4 in FGF21-deficient mice abolishes the anti-atrophic effects, indicating that cytoplasmic HDAC4 is a key mediator of FGF21 deficiency-derived protective effects. Additionally, Class IIa HDACs such as HDAC5 and HDAC7 have been reported to function compensatorily in the liver to regulate glucose hemostasis (61). We observed concurrent phosphorylation and cytoplasmic accumulation of HDAC5 and HDAC7 in the absence of FGF21, which may also contribute to the anti-atrophic effects following denervation (data now shown). Moreover, HDAC4, as an enzyme, is supposed to interact with cytoplasmic substrates to enhance NMJ innervation. Future studies are needed to explore the potential compensatory effects of these HDACs and their associated substrates in response to neurogenic muscle atrophy.

TGFB1 triggers diverse effects by stimulating TGFB receptors in conjunction with downstream transcription factors or kinases (62). While TGFB1’s role in promoting denervation-induced muscle atrophy is well-documented (63, 64), its interaction with FGF21 in this context has not been studied. Our findings reveal that the skeletal muscle-specific expression pattern of TGFB1 upon denervation, which mirrors that of FGF21, strongly supports its local induction of FGF21, initiating downstream events.

We further demonstrate that TGFB1 induces FGF21 expression in differentiated muscle cells via the non-canonical JNK/c-Jun axis, leading to NMJ instability and eventually muscle atrophy. The presence of augmented TGFB signaling in ALS (65), coupled with FGF21’s ability to impair NMJs, raises the question of whether the TGFB1/FGF21 axis represents a generalized signaling pathway in neurodegenerative conditions, warranting further investigation. TGFB1, a known secreted cytokine, must originate from an intramuscular cellular source activated by denervation, contributing to muscle atrophy. Madaro et al. reveal that FAPs extracted exclusively from denervated muscle, as opposed to injured muscle, are capable of inducing muscle atrophy (66). Kajabadi et al. report that cachectic muscle FAPs produce Activin-A, a member of the TGFB superfamily, to induce muscle atrophy (67). Based on these findings, we speculate that denervated FAPs might secrete TGFB1 to drive muscle atrophy. In our study, we observed greatly higher TGFB1 expression in denervated FAPs than in non-denervated FAPs, which effectively promote muscle atrophy. FAPs can be activated to form specific subpopulations that expand and adopt different fates (68). Our findings show that 14-day denervated FAPs adopt a pro-atrophic ability, initiating downstream TGFB1-mediated pro-atrophic programs. This 14-day period corresponds well with the pro-atrophic effect of FAPs, as neither injured FAPs nor 3-day denervated FAPs exhibit this effect (66), indicating that severe NMJ loss over a specific period is critical for FAPs activation. Furthermore, the expression of the FAPs marker, PDGFRα is enhanced along with TGFB1 in ALS muscle, suggesting that FAPs-derived TGFB signaling could be a promising target to counteract muscle atrophy following denervation or during ALS progression.

To our knowledge, this is the first study to demonstrate that FGF21 is a key factor that regulates neurogenic skeletal muscle atrophy. Using both gain-of-function and loss-of-function approaches, we elucidate a novel role for FGF21 in regulating skeletal muscle mass through its effects on NMJ innervation. These findings are critical for understanding the molecular pathways governing muscle mass and highlight the need to consider the potential adverse effects of FGF21 on specific organs when developing FGF21 agonists for treating metabolic disorders.

## Methods

### Sex as a biological variable

All experiments involving animals were conducted using male mice at 5 to 11 weeks of age. It is unknown whether the findings are relevant for female mice.

### Animal studies

All mice used in our experiments were housed in pathogen-free conditions under a standard 12:12-h light/dark cycle, with ad libitum access to a standard chow diet (Labo MR Stock, Nosan Corporation Bio-Department) and water. Wilde-type C57BL/6J mice (WT), mdx mice (C57BL/10ScSn-Dmd^mdx^/J) and control mice (C57BL/10ScSn) were purchased from Crea-Japan Inc. Aged mice (24 months old) and young mice (3 months old) were generously provided by Dr. Ishigami (Tokyo Metropolitan Institute for Geriatrics and Gerontology, Japan). Fgf21 global knockout (Fgf21KO) mice were kindly provided by Dr. Nobuyuki Itoh (Kyoto University, Japan). The genotypes of mice were determined as previously described (69).

### Animal models of muscle atrophy

Several muscle atrophy models were employed, including starvation, denervation, muscular dystrophy, and aging. In the starvation experiments, animals were deprived of food for 24 or 48 hours before being sacrificed. For denervation, one limb of the mouse was shaved and sterilized, the sciatic nerve was exposed, and a 5-10 mm segment was excised. The incision was then sutured using an 8-0 sterile silk suture. The contralateral limb, subjected to the same operation without sciatic nerve transection, served as the sham group. Different denervation strategies were selected based on experimental objectives. For general biological experiments (e.g., gene expression, protein expression, and histological analysis) and in vivo Hdac4 knockdown studies, unilateral sham surgery and contralateral denervation were performed on the same mouse. For grip strength tests and evaluation of Fgf21 expression across different tissues, both limbs of a mouse were either sham-operated or denervated separately. In experiments involving in vivo Fgf21 overexpression or ELISA assays of plasma, one limb of a mouse was either sham-operated or denervated, respectively. Mice were sacrificed at 3-, 7-, 14-or 31-days post-surgery. The muscular dystrophy and aging models directly utilized mdx mice and aged mice, respectively. Muscles collected at the specified times were either prepared for histological analysis or frozen in liquid nitrogen and stored at-80°C for future use.

### In vivo electroporation

Eleven-week-old male C57BL/6J mice were used for in vivo electroporation experiments. The day before electroporation, hair was removed from the hindlimbs. On the day of the procedure, mice were anesthetized, and 100 μL of hyaluronidase (40 U/mL) was injected into tibialis anterior (TA) muscle. After a 2-hour wait, 30 μL of plasmid solution was injected into both TA muscles. Electroporation was performed using the NEPA21 Super Electroporator with CUY560-3-0.5 electrodes, measuring the impedance and applying voltage with a 1-second pulse interval for a total of 3 pulses. For starvation experiments, 1 μg/μL of pEF2-βgal or pEF2-FGF21 was electroporated 10 days before the 48-hour starvation period.

### AAV constructs, production, and purification

The AAV2/8-TBG-GFP expression plasmid, p5E18-VD2/8 helper plasmid, and pXX6-80 capsid plasmid were generously provided by Dr. Itoh (70). AAV serotype 8 is known for its proven tropism for skeletal muscle cells (71). To drive in vivo muscle-specific gene expression, the 1,350 base pair region of the pBS MCK promoter (Addgene, #12528), which has been shown to efficiently and specifically drive gene expression in skeletal muscle (72), was amplified and used to replace the TBG promoter of AAV2/8-TBG-GFP using PacI/EcoRI restriction enzyme sites, resulting in the AAV2/8-MCK-GFP plasmid. To achieve in vivo gene knockdown, the GFP-U6 expression cassettes were excised from the AAV2/1-GFP-U6 plasmid, a gift from Dr. Eguchi (73), and used to replace the TBG promoter and GFP of AAV2/8-TBG-GFP using PacI/HindIII restriction enzyme sites, generating the AAV2/8-GFP-U6 plasmid. To overexpress Fgf21, the coding sequence for mouse Fgf21 was inserted to replace the GFP sequence in AAV2/8-MCK-GFP, with the AAV2/8-MCK-GFP serving as a control virus. To silence Hdac4 expression, two shRNA sequences targeting mouse Hdac4 (61, 74) or a non-targeting control sequence (75) were inserted into the AAV2/8-GFP-U6 plasmid (resulting in AAV2/8-GFP-shHdac4 and AAV2/8-GFP-shCtrl, respectively). These constructs were placed under the control of the U6 promoter, with GFP serving as a reporter to monitor shRNA delivery efficiency. AAV vectors used for overexpression or knockdown experiments were packaged by triple transfection of expression plasmids, helper plasmids, and capsid plasmids into HEK293T cells, followed by purification via CsCl density gradient ultracentrifugation (76). Viral titers were determined using TaqMan real-time quantitative PCR.

### AAV injections in skeletal muscles

For overexpression experiments, 5-week-old male C57BL/6J mice were intramuscularly injected with 2.5 × 10¹⁰ viral genomes (vg) of either AAV2/8-MCK-GFP or AAV2/8-MCK-mFGF21 vectors into the tibialis anterior (TA) muscles. For knockdown experiments, 5-week-old male C57BL/6J mice were intramuscularly injected with 1 × 10¹¹ vg of either AAV2/8-GFP-shCtrl or AAV2/8-GFP-shHdac4 vectors into TA muscles. Four weeks post-injection, denervation surgery was performed as previously described.

### Histological analysis

TA muscle samples were snap-frozen in liquid nitrogen-precooled isopentane, embedded in Tissue-Tek O.C.T. compound (Sakura Finetek, Japan), and sectioned into 10 μm cryosections. For Hematoxylin and Eosin (H&E) staining, cryosections were air-dried, fixed in 4% paraformaldehyde, washed in PBS, and stained with hematoxylin and eosin following standard protocols. The cross-sectional area was measured using Fiji software, as described in a published protocol (77).

### Grip strength

A grip strength meter (Muromachi, MK-380V) was used to measure grip strength. Mice were allowed to grasp the device’s metal grid with all four paws and were gently pulled backward by their tails in a straight line until the grip was broken and the peak force measurement was recorded. Each mouse was tested five times with 30-second intervals between tests. The highest and lowest values were excluded, and the remaining three force measurements were averaged and normalized to body weight.

### Subcellular fractionation

Nuclear protein extraction from tissues was performed according to a previously published protocol (78). Briefly, 50 mg of TA muscles were homogenized in STM buffer (250 mM sucrose, 50 mM Tris–HCl pH 7.4, 5 mM MgCl2), and the supernatant was precipitated using four times the volume of pre-chilled acetone. After acetone was completely removed, the resulting pellet was resuspended and used as the cytosolic fraction. For the nuclear fraction, the pellet obtained in the first step was washed three times in STM buffer and then resuspended in NET buffer (20 mM HEPES pH 7.9, 1.5 mM MgCl2, 0.5 M NaCl, 0.2 mM EDTA, 20% glycerol, 1% Triton X-100). The suspension was then passed through an 18-gauge needle, and the supernatant was collected as the nuclear fraction. All procedures were performed on ice, and protease and phosphatase inhibitor cocktails were added to each buffer immediately before use.

### Whole-mount neuromuscular junction (NMJ) staining

The protocol for NMJ staining of extensor digitorum longus (EDL) muscles was adapted from a published method (79). EDL muscles were dissected and fixed in 4% paraformaldehyde (PFA) in PBS, then dehydrated in 30% sucrose. After rinsing with PBS, muscles were incubated in 0.1 M glycine to quench formaldehyde-induced autofluorescence. The muscles were then separated into four pieces and permeabilized in a solution containing 0.5% Triton X-100 and 4% BSA, followed by incubation with primary antibodies: mouse anti-NF-M (1:200, 2H3, DSHB) and mouse anti-SV2 (1:200, 2H3, DSHB) in a blocking solution containing 2% Triton X-100 and 4% BSA. After washing with 2% Triton X-100 in PBS, the muscles were incubated with secondary antibody Alexa Fluor™ 488 (1:500, A11029, Invitrogen) and α-BTX (1:500, Invitrogen) for 2 hours. The muscles were then washed again with 2% Triton X-100 in PBS, mounted with ProLong™ Glass Antifade with NucBlue™ (Thermo Fisher Scientific), flattened with magnets, and sealed with nail polish. Samples were stored in the dark until imaging with a ZEISS LSM 800 confocal laser scanning microscope (Carl Zeiss).

### Cell isolation

Murine primary cells from denervated mice were isolated using magnetic-activated cell sorting (MACS, Miltenyi). Briefly, all hindlimb skeletal muscles were dissected, mechanically dissociated, and digested in an enzymatic mix containing 2 mg/mL collagenase D, 2.4 U/mL dispase II, and 10 μg/mL DNase I, supplemented with 1 mM CaCl2 and 5 mM MgCl2. The digested muscles were then passed through an 18-gauge needle several times. The resulting mixture was sequentially filtered through 100, 70, and 40 μm cell strainers and incubated with RBC lysis buffer to remove erythrocytes. Different cell populations were then isolated through consecutive incubation with microbead-conjugated antibodies in the cell suspension. Lin+/CD31+ cells were collected as lineage cells and endothelial cells. Satellite cells (SCs) were identified as Lin-/CD31-/α7-integrin+ cells, while fibro/adipogenic progenitors (FAPs) were defined as Lin-/CD31-/α7-integrin-, PDGFRα+ cells.

### SCs and FAPs culture and conditioned medium (CM) collection

Freshly isolated FAPs were cultured in Dulbecco’s modified Eagle’s medium (DMEM) containing 20% FBS until they reached confluence. The cells were then washed with PBS and cultured in serum-free DMEM for 24 hours. The conditioned medium (CM) was collected and centrifuged at 3,000 rpm for 10 minutes at 4°C, and the supernatant was stored at-80°C until use. CM treatments were applied to 3-day differentiated C2C12 cells for the specified periods.

Freshly isolated satellite cells were cultured in DMEM containing 10% FBS, 10% horse serum, and 0.5% chick embryo extract. To induce differentiation, the medium was switched to DMEM containing 5% horse serum. Undifferentiated satellite cells and 4-day-differentiated satellite cells were treated with 5 ng/mL TGFB1 for 4 hours to assess Fgf21 mRNA expression.

### RNA sequencing (RNA-seq) and public datasets analysis

Two weeks post-denervation, TA muscles from sham-operated or denervated mice (n=4 per group) were collected. Total RNA was extracted using the RNeasy kit (QIAGEN), and the RNA from each group was pooled and submitted to Macrogen Co., Ltd for analysis. Differentially expressed genes (DEGs) were analyzed using the edgeR software package, with DEGs identified based on an adjusted p-value < 0.05 and |logFC| ≥ 1. Data analysis and visualization were performed in R (v 4.3.1). Upregulated genes used to generate a Venn diagram were obtained from the NCBI Gene Expression Omnibus (GEO) datasets: GSE18119 (mouse quadriceps muscle, mitochondrial myopathy) (unpublished), GSE48574 (human skeletal muscle biopsies, iron-sulfur [Fe-S] cluster-deficient myopathy) (80), GSE52766 (mouse gastrocnemius muscle, mdx) (81), GSE49826 (mouse TA muscle, 2-week denervation) (82), and GSE87108 (mouse gastrocnemius muscle, 28-month-old) (83). The screening threshold was set at p < 0.05. Additionally, the expression of Fgf21 in denervated TA muscles of rats was analyzed using GSE201025 (rat TA muscle, 2-week denervation) (84).

### Plasmid constructs

The coding sequence of mouse Hdac4 was cloned into a p3XFLAG-CMV-7.1 vector (Sigma-Aldrich) using HindIII and EcoRI restriction enzyme sites to construct the FLAG-Hdac4 expression plasmid.

### Cell culture

Murine myoblasts C2C12 (obtained from ATCC) were maintained at 37°C and 5% CO2 in high glucose DMEM supplemented with 10% FBS. Cells were sub-cultured when they reached 50-60% confluence. Once confluent, the culture medium was replaced with a differentiation medium, consisting of high glucose DMEM supplemented with 2% horse serum, to induce myotube formation. Fresh differentiation medium was replenished every 2 days, and cells were allowed to differentiate for 3-4 days before experiments. The passage number of C2C12 cells used in this research did not exceed 20.

### RNA interference

siRNAs targeting Fgf21, Smad3, and control siRNA were purchased from Dharmacon (ON-TARGETplus Mouse Fgf21 [56636] siRNA–SMARTpool, ON-TARGETplus Mouse Smad3 [17127] siRNA–SMARTpool, and ON-TARGETplus Non-targeting pool). C2C12 cells that had been differentiated for 2 days were transfected with control siRNA (10 or 30 nM), or siRNA targeting mouse Fgf21 (10 nM) or Smad3 (30 nM) using Lipofectamine RNAiMAX Transfection Reagent (Thermo Fisher) according to the manufacturer’s instructions. 24 or 48 hours after transfection, cells were used for the indicated experiments as detailed in the results section.

### Immunofluorescence (IF) staining

Myotube diameter and the localization of exogenous or endogenous HDAC4 were determined by confocal microscopy following a standard immunofluorescence (IF) staining protocol. C2C12 cells were washed, fixed with 4% paraformaldehyde (PFA), permeabilized with 0.5% Triton X-100, and blocked with 3% BSA. The specimens were then incubated with anti-MyHC (MF20; R&D Systems), anti-HDAC4 (CST), or anti-FLAG (Sigma) antibodies for 2 hours, followed by incubation with Alexa Fluor 488–conjugated goat anti-mouse or anti-rabbit IgG antibodies (1:500) for 1 hour. The specimens were mounted with ProLong™ Glass Antifade with NucBlue™ (Thermo Fisher Scientific) and observed using a ZEISS LSM800 confocal laser scanning microscope (Carl Zeiss).

### Acetylcholine receptor (AChR) Clustering Assay

The stability of AChR clusters was assessed as previously described (85). Briefly, after 4 days of differentiation to induce AChR clusters, C2C12 myotubes were treated with Agrin alone or in combination with other compounds for 16-18 hours and then maintained in a medium with or without additional compounds. Cells were fixed in 2% PFA, and AChRs were stained with α-bungarotoxin 594 (BTX; 1 mg/mL, Invitrogen) diluted 1:2500. After washing, cells were mounted with ProLong™ Glass Antifade with NucBlue™ (Thermo Fisher Scientific). Samples were stored and protected from light until imaging with a ZEISS LSM 800 confocal laser scanning microscope (Carl Zeiss).

### Immunoblotting

C2C12 cells and mouse tissues were lysed in RIPA buffer containing PMSF (Sigma-Aldrich, MO), a protease inhibitor cocktail (Nacalai Tesque), and a phosphatase inhibitor cocktail (Sigma-Aldrich). The protein lysates were then boiled and subjected to SDS-PAGE, followed by incubation with primary and secondary antibodies according to standard procedures. Antibodies used are shown in **Supplementary Table 1**.

### Quantitative real-time PCR (RT-qPCR)

Total RNA from cells or mouse tissues was extracted using ISOGEN (NIPPON GENE) according to the manufacturer’s instructions and reverse transcribed into cDNA using the High-Capacity cDNA Reverse Transcription Kit (Applied Biosystems). Quantitative real-time PCR was performed on an Applied Biosystems StepOnePlus instrument using SYBR Green Master Mix (Thermo Fisher Scientific). qPCR primers used are shown in **Supplementary Table 2**.

### Enzyme-linked immunosorbent assay (ELISA)

The protein levels of FGF21 and TGFB1 in plasma, muscle tissue, and culture medium were measured using a Mouse/Rat FGF-21 ELISA kit (R&D Systems) or a Human/Mouse/Rat/Porcine/Canine TGF-beta 1 ELISA kit (R&D Systems), according to the manufacturer’s instructions.

### Statistical analysis

All data are presented as mean ± SD. An unpaired two-tailed Student’s *t*-test was used for comparisons between the two groups. For comparisons involving more than two groups, one-way analysis of variance (ANOVA) followed by the Tukey–Kramer post hoc test was applied. Statistical significance is denoted in the figures as *(p < 0.05), **(p < 0.01), and ***(p < 0.001). Statistical analysis was performed using GraphPad Prism 9.0 software.

### Study approval

All animal studies were conducted in accordance with the guidelines approved by the Animal Usage Committee of the University of Tokyo (Law No. 105, 1 October 1973, as amended on 1 June 2020).

## Author contributions

Conceptualization: LZ, TS, MS, and RS. Investigation: LZ, TS, LN, YT, YY and MS. Funding acquisition: MS, RS. Supervision: TS, MS, and RS. Writing – original draft: LZ. Writing – review and editing: TS, and RS.

## Acknowledgments

We are grateful to Dr. Nobuyuki Itoh (Medical Innovation Center, Graduate School of Medicine, Kyoto University, Kyoto) for providing the whole body Fgf21-knockout mice. This research was supported by grants from JSPS KAKENHI Grant-in-Aid for Scientific Research (B): 19H02906 (to MS), JSPS KAKENHI Grant-in-Aid for Exploratory Research: 21K19066 (to MS) and JSPS KAKENHI Grant-in-Aid for Scientific Research (A): 20H00408 (to RS).

